# Robust associations between white matter microstructure and general intelligence

**DOI:** 10.1101/2022.05.02.490274

**Authors:** Christina Stammen, Christoph Fraenz, Rachael G. Grazioplene, Caroline Schlüter, Viola Merhof, Wendy Johnson, Onur Güntürkün, Colin G. DeYoung, Erhan Genç

**Author notes:** Corresponding author: Dr. Erhan Genç Address: Department of Psychology and Neuroscience, Leibniz Research Centre for Working Environment and Human Factors (IfADo), Ardeystraße 67, 44139 Dortmund, Germany.

## Abstract

Early research on the neural correlates of human intelligence was almost exclusively focused on gray matter properties. The advent of diffusion-weighted imaging led to an exponential growth of white matter brain imaging studies. However, this line of research has yielded mixed observations, especially about the relations between general intelligence and white matter microstructure. We used a multi-center approach to identify white matter regions that show replicable structure-function associations, employing data from four independent samples comprising over 2000 healthy participants. We used tract-based spatial statistics to examine associations between *g* factor scores and white matter microstructure and identified 188 voxels which exhibited positive associations between *g* factor scores and fractional anisotropy in all four data sets. Replicable voxels formed three clusters: one located around the forceps minor, crossing with extensions of the anterior thalamic radiation, the cingulum-cingulate gyrus, and the inferior fronto-occipital fasciculus in the left hemisphere, one located around the left-hemispheric superior longitudinal fasciculus, and one located around the left-hemispheric cingulum-cingulate gyrus, crossing with extensions of the anterior thalamic radiation and the inferior fronto-occipital fasciculus. Our results indicate that individual differences in general intelligence are robustly associated with white matter organization in specific fiber bundles.

---

People differ in general intelligence, i.e. “[…] their ability to understand complex ideas, to adapt effectively to the environment, to learn from experience, to engage in various forms of reasoning, to overcome obstacles by taking thought” (Neisser et al. 1996). As discovered by Spearman (1904), individuals who do well in one cognitive task tend to perform above average in other cognitive tasks as well. The phenomenon of positively correlated cognitive test scores, which he termed the ‘positive manifold’, led Spearman to declare the existence of ‘*g*’, the general factor of intelligence. Though *g* is actually just a statistical observation, it is an important one because it is relevant to many aspects of everyday life. For example, intelligence is positively correlated with school performance (Neisser et al. 1996; Roth et al. 2015), job performance (Gottfredson 1997; Schmidt and Hunter 2004), socioeconomic success (Strenze 2007), income (Zagorsky 2007), and even physical health, longevity, and ephemerals such as stability of marital relationships (Aspara et al. 2018; Batty et al. 2007; Calvin et al. 2017; Calvin et al. 2011; Deary et al. 2010b; Hemmingsson et al. 2006; Whalley and Deary 2001). Due to the impacts that intelligence or *g* seems to have on life outcomes, it has always been of interest to identify specific structures within the human brain that are associated with its interindividual differences. Although different individuals use different strategies to solve any particular intelligence test task and that the same individual uses different strategies at various times, is it possible to find general patterns that show up consistently when doing analyses across many individuals.

While it is one well-replicated observation that bigger brains are weakly to moderately associated with higher intelligence (Cox et al. 2019; McDaniel 2005; Pietschnig et al. 2015), the advent of *in vivo* neuroimaging techniques has allowed scientists to move from overall brain size to various properties of single brain regions. Jung and Haier (2007) reviewed 37 neuroimaging studies that aimed to identify intelligence-related brain regions using various intelligence measures and imaging techniques. Based on the commonalities across findings, they proposed the Parieto-Frontal Integration Theory (P-FIT) of intelligence. P-FIT nominates a set of distributed brain regions, mainly located in parietal and frontal areas, whose functional and structural properties are related to interindividual intelligence differences. Each of these P-FIT areas is believed to be involved in one of the multiple information processing stages used in solving abstract reasoning tasks. Hence, efficient and flawless information transfer between these regions is considered to play vital roles in intellectual achievements, which in turn indicates roles of brain white matter (Jung and Haier 2007). The brain’s white matter mainly consists of myelinated axons that are organized in fiber tracts running from one brain region to another (Filley 2012), which enables thereby the information transfer. The hypothesis that the integrity of certain white matter fiber tracts is crucial for intelligence has been empirically supported by Gläscher et al. (2010) who used voxel-based lesion-symptom mapping in a large sample of patients with focal brain damage. Their observations indicated that severe damage to fiber tracts linking P-FIT areas was significantly associated with lower intelligence. More precisely, they observed negative associations in tracts connecting fronto-parietal areas (superior longitudinal fasciculus, arcuate fasciculus), fronto-temporal areas (uncinate fasciculus), and fronto-occipital areas (fronto-occipital fasciculus) (Gläscher et al. 2010). Subsequent studies using lesion-symptom mapping were consistent with these observations (Barbey et al. 2014; Barbey et al. 2012; Bowren et al. 2020).

The advent of diffusion-weighted imaging (DWI) led to an exponential growth of white matter brain imaging studies (Deary et al. 2022). DWI is based on diffusion of water molecules (Le Bihan 2014; Le Bihan and Breton 1985; Le Bihan et al. 1986) and provides opportunity to study white matter microstructure *in vivo* and non-invasively (Tournier et al. 2011). Since white matter mainly consists of myelinated axons (Filley 2012), axon membranes form natural borders for water molecules and limit their movements perpendicular to the fibers (Le Bihan 2003). Therefore, DWI indicates anisotropic, directional diffusion patterns within voxels containing coherently oriented white matter fibers and isotropic, non-directional patterns within voxels containing randomly oriented fibers or fluid-filled spaces such as ventricles (Le Bihan 2003). The most widely used metric to quantify water diffusion’s degrees of directionality in a summative manner is fractional anisotropy (FA). Here, higher FA values indicate more parallel diffusion trajectories (Assaf and Pasternak 2008; Basser and Pierpaoli 1996). Although FA is clearly related to white matter microstructure, it may be misleading to use it as a marker of microstructural integrity within specific fiber bundles because lower FA values do not necessarily indicate some kind of tissue damage (Jones et al. 2013). FA is a complex and unspecific measure affected by various physiological factors like axon diameter, fiber density, myelin concentration, or distribution of fiber orientation (Beaulieu 2002; Friedrich et al. 2020; Jones et al. 2013; Le Bihan 2003). The latter refers to the ‘crossing fiber problem’, which describes that FA values are inaccurate in voxels with complex fiber architectures such as multiple fiber populations, bending fibers, or crossing fibers (Jeurissen et al. 2013; Jones et al. 2013). These various physiological factors make it challenging to disentangle and interpret the actual sources of signal differences (Jones et al. 2013). Nevertheless, FA is a widely used metric and its association with intelligence has been investigated extensively. Studies have analyzed white matter properties by averaging across specific regions of interest (Deary et al. 2006; Power et al. 2019; Tang et al. 2010), extracting them from whole fiber tracts (Bathelt et al. 2019; Booth et al. 2013; Clayden et al. 2012; Cox et al. 2019; Cremers et al. 2016; Dubner et al. 2019; Ferrer et al. 2013; Fuhrmann et al. 2020; Gongora et al. 2020; Holleran et al. 2020; Kennedy et al. 2021; Kievit et al. 2016; Kievit et al. 2014; Kievit et al. 2018; Kontis et al. 2009; Muetzel et al. 2015; Nestor et al. 2015; Ohtani et al. 2014; Penke et al. 2012; Penke et al. 2010; Simpson-Kent et al. 2020; Suprano et al. 2020; Urger et al. 2015; Yu et al. 2008), or by a whole-brain voxel-based approach (Allin et al. 2011; Chiang et al. 2009; Navas-Sanchez et al. 2014; Schmithorst 2009; Schmithorst et al. 2005). As summarized by Genç and Fraenz (2021), the majority of such studies reported positive relations between intelligence and average FA values from many major white matter pathways, mostly representing connections between P-FIT areas. Independent of the specific methods used, similar patterns emerged among different studies. The four fiber tracts most commonly associated with intelligence differences are the genu and the splenium of the corpus callosum, the uncinate fasciculus, and the superior longitudinal fasciculus (Genç and Fraenz 2021).

Studies investigating pre-selected brain regions or white matter tracts are prone to miss relevant relations in non-selected areas. Analyses adapting voxel-based methods, such as voxel-based morphometry (Ashburner and Friston 2000), to analyze FA images also have various shortcomings such as alignment inaccuracies or the dependence of the results on the arbitrarily chosen extent of spatial smoothing (Smith et al. 2006). Tract-Based Spatial Statistics (TBSS) has been introduced as an approach that combines the strengths of tractography-based and voxel-based analyses to overcome the aforementioned limitations (Smith et al. 2006). Although TBSS has advantages, few studies have investigated the relation between FA and intelligence in healthy (young) adults using this method. Dunst et al. (2014) found no significant associations between general intelligence and FA in any white matter voxel, whereas Malpas et al. (2016) reported significant positive relations in 32% of voxels constituting the white matter skeleton (right anterior thalamic radiation, left superior longitudinal fasciculus, left inferior fronto-occipital fasciculus, and left uncinate fasciculus). In line with Dunst et al. (2014), Hidese et al. (2020) found no significant associations between general intelligence and regional white matter FA, despite analyzing a larger sample. However, FA showed significant positive relations with scores from specific subtests, namely processing speed and symbol search (Hidese et al. 2020). Tamnes et al. (2010) employed a sample comprised of 168 participants, aged between 8 and 30 years. While they focused their TBSS analyses on verbal and performance abilities, they also conducted a tract-based approach for full-scale intellectual abilities and observed significant positive relations between general intelligence and FA values in almost all analyzed fiber tracts. The TBSS analyses revealed significant positive associations between FA and verbal abilities in 4.6% of voxels in the white matter skeleton (left anterior thalamic radiation, left cingulum-cingulate gyrus, left and right superior longitudinal fasciculus). Performance abilities were significantly associated with FA in 1.6% of skeleton voxels (left superior longitudinal fasciculus, forceps major; Tamnes et al. 2010).

Previous TBSS studies have often had samples small enough that effect size estimates are likely to be highly variable and inaccurate. Furthermore, inconsistencies such as different sample sizes or intelligence measures limited their comparability. In short, they do not allow clear conclusions to be drawn about associations between general intelligence and FA. Some found significant positive relations while others did not. As proposed by Genç and Fraenz (2021), such inconsistent findings may be tackled by following a multi-center approach. To this end, multiple data sets, typically collected by different research groups, are analyzed independently. Importantly, only those results which replicate across the majority (or all) of samples are considered robust. We followed this approach methodologically consistently as possible, searching for replicable observations among four independent data sets comprising cross-sectional data from more than 2000 healthy participants. We performed whole-brain TBSS analyses to examine the associations between general intelligence, in the form of *g* factor scores, and FA separately on each sample. Besides the aforementioned advantage of multi-center studies, another reason for choosing this rather conservative approach was that the combination of our four data sets was not possible since individuals’ *g* levels in different samples might differ from each other and because imaging data were obtained on different scanners. Data were collected at Ruhr-University Bochum (RUB) in Germany with N = 557 (Genç et al. 2021), the Human Connectome Project (HCP) with N = 1206 (van Essen et al. 2013), the University of Minnesota (UMN) with N = 335 (Grazioplene et al. 2016; Grazioplene et al. 2015), and the Nathan Kline Institute (NKI) with N = 417 (Nooner et al. 2012). We compared observations to identify white matter areas that exhibited replicable structure-function associations among data sets.

## Materials and Methods

### Data set RUB

#### Participants

The RUB sample encompassed 557 participants, mainly university students of different majors, who were either paid for their participation or received course credits. Although the age range was between 18 and 75 years, the data set was predominantly comprised of individuals from young adulthood (mean age: 27.3 years, SD = 9.4 years). There were 283 males and 274 females. Individuals were not admitted to the study if they had insufficient German language skills or reported having undergone any of the employed intelligence tests within the last five years. They were also excluded if they or any of their close relatives suffered from neurological and/or mental illnesses, as assessed by a self-report questionnaire. The study protocol was approved by the local ethics committee of the Faculty of Psychology at Ruhr University Bochum (vote Nr. 165). All participants gave written informed consent and were treated according to the Declaration of Helsinki.

#### Measurement of intelligence

##### I-S-T 2000 R

The Intelligenz-Struktur-Test 2000 R (I-S-T 2000 R; Liepmann et al. 2007) is a broadly applicable, well-established German intelligence test battery that takes about 2.5 hours to complete. It measures multiple intelligence facets as well as general intelligence. Most included cognitive tasks are presented in multiple-choice format. Verbal, numerical, and figural abilities are each assessed by three different mental reasoning tasks of 20 items. Verbal intelligence is assessed by tasks in which participants must complete sentences (IST_SEN), find analogies (IST_ANA), and recognize similarities (IST_SIM). Numerical intelligence is assessed by tasks involving arithmetic calculations (IST_CAL), number series (IST_SER), and mathematical equations to which arithmetic signs need to be added (IST_SIG). Figural intelligence is assessed by tasks in which participants must select and reassemble parts of a cut-up figure (IST_SEL), mentally rotate and match three-dimensional objects (IST_CUB), and solve matrix-reasoning problems (IST_MAT). In addition, retention (IST_RET) is assessed by 23 verbal and figural items. Here, participants must memorize series of words or figure pairs. An extension module comprises 84 multiple-choice questions on six knowledge facets (art/literature, economy, geography/history, mathematics, science, and daily life) and measures general knowledge (IST_KNO). Reliability estimates (Cronbach’s *α*) are between .88 and .96 for subtests and composite scores (Liepmann et al. 2007).

##### BOMAT-Advanced Short

The Bochumer Matrizentest (BOMAT; Hossiep et al. 2001) is a non-verbal German intelligence test whose structure is comparable to the well-established Raven’s Advanced Progressive Matrices (Raven et al. 1990). For the study at hand, we used the advanced short version, which is widely used in neuroscientific research and known to have high discriminatory power in samples with generally high intellectual abilities, thus avoiding possible ceiling effects (Fraenz et al. 2021; Genç et al. 2018; Genç et al. 2019; Hossiep et al. 2001; Oelhafen et al. 2013). The test comprises 29 matrix-reasoning items. Each item shows a 5-by-3 matrix composed of elements arranged according to a specific but unspecified rule. One field within the matrix is empty and needs to be filled with of six provided elements that follows the rule. Split-half reliability of the BOMAT is .89 and Cronbach’s *α* is .92 (Hossiep et al. 2001).

##### BOWIT

The Bochumer Wissenstest (BOWIT; Hossiep and Schulte 2008) is a German general knowledge questionnaire. It assesses eleven different knowledge facets, from two major domains. The four facets biology/chemistry, mathematics/physics, nutrition/exercise/health, and technology/electronics are assigned to the scientific-technical knowledge domain. The social and humanistic knowledge domain has seven facets: arts/architecture, civics/politics, economics/laws, geography/logistics, history/archeology, language/literature, and philosophy/religion. The BOWIT is available in two parallel test forms, in which each knowledge facet is represented by 14 multiple-choice questions. To measure general knowledge as precisely as possible, all participants had to complete both test forms, resulting in 308 items. The BOWIT shows reliability estimates greater than .90: split-half reliability is reported as .96, Cronbach’s *α* .95, test-retest reliability .96, and parallel-form reliability .91 (Hossiep and Schulte 2008).

##### ZVT

The Zahlenverbindungstest (ZVT; Oswald and Roth 1987) is a trail-making test to assess the cognitive processing speed of both children and adults. The test consists of two short sample tasks and four assessed tasks. Here, participants connect numbers from 1 to 90 based on a specific rule as fast as possible. The processing times for the four tasks are averaged to obtain a comprehensive measure of processing speed. The reliability across the four tasks is reported as .95 in adults. The six-month retest-reliability is reported to be between .84 and .90 (Oswald and Roth 1987).

##### Acquisition of diffusion-weighted imaging data

All images were collected on a Philips 3T Achieva scanner at Bergmannsheil Hospital in Bochum, Germany, using a 32-channel head coil. Diffusion-weighted images were acquired using echo planar imaging (TR = 7652 ms, TE = 87 ms, flip angle = 90°, 60 slices, matrix size = 112 x 112, voxel size = 2 x 2 x 2 mm). Diffusion weighting was uniformly distributed along 60 directions using a *b*-value of 1000 s/mm^2^. Additionally, six volumes with no diffusion weighting (*b* = 0 s/mm^2^) were acquired as an anatomical reference for motion correction. To increase the signal-to-noise ratio of diffusion-weighted images, we acquired three consecutive scans that were subsequently averaged (Genç et al. 2011a; Genç et al. 2011b). The total acquisition time was 30 minutes.

### Data set HCP

#### Participants

Data were provided by the Human Connectome Project, WU-Minn Consortium (Principal Investigators: David Van Essen and Kamil Ugurbil; 1U54MH091657), funded by the 16 United States National Institutes of Health (NIH) Institutes and Centers supporting the NIH Blueprint for Neuroscience Research and by the McDonnell Center for Systems Neuroscience at Washington University. We employed the “1200 Subjects Data Release” (van Essen et al. 2013). It includes behavioral and imaging data from 1206 young adults. To compute a *g* factor, all participants with missing values in one or more of the intelligence measurements listed below had to be excluded, which reduced the sample to N = 1188 (mean age: 28.8 years, SD = 3.7 years, 641 females). Since DWI data were not available for all participants, the final sample for the TBSS analysis was limited to 1061 participants (mean age: 28.7 years, SD = 3.7 years, 571 females). The age range was between 22 and 37 years. To be included in the data set, participants had to have no significant history of psychiatric disorder, substance abuse, neurological, or cardiological disease and give valid informed consent (van Essen et al. 2012).

#### Measurement of intelligence

##### Penn CNB

Four subtests from the University of Pennsylvania Computerized Neurocognitive Battery (PennCNB; Gur et al. 2001; Gur et al. 2010; Moore et al. 2015) were used to assess intelligence. These included the Penn Matrix Reasoning Task (PMAT), the Short Penn Continuous Performance Test (SCPT), the Variable Short Penn Line Orientation Test (VSPLOT), and the Penn Word Memory Test (IWRD).

The PMAT is an abbreviated version of Raven’s Standard Progressive Matrices Form A (Bilker et al. 2012) and measures nonverbal reasoning ability. In 24 items, responders infer which of five options most reasonably belongs in the one empty square of a 2-by-2, 3-by-3 or 1-by-5 matrix. Items are presented with increasing difficulty and the task discontinues if the participant selects incorrect answers to five consecutive items. The SCPT is a measure of visual attention. Lines are presented on a computer screen for 300 milliseconds, followed by a black screen for 700 milliseconds. Participants are supposed to indicate when the lines form a number (in the first experiment block) or a letter (in the second experiment block). Each block consists of 90 stimuli. The VSPLOT is used to assess visual-spatial processing.

Participants see two lines on a computer screen and their task is to rotate one line so that it is parallel to the other. The VSPLOT comprises 24 trials. The IWRD (Form A) measures verbal episodic memory. Participants are shown 20 words, each for one second, which they are supposed to memorize. During the recognition phase, 40 words are presented - the previously presented 20 words and 20 distractors which are matched for length, imageability, and concreteness. Participants indicate whether each of the 40 words was one of the memorized 20 words (Gur et al. 2001; Gur et al. 2010; Moore et al. 2015).

The reliability estimates (Cronbach’s *α*) for all subtests of the Penn CNB are reported to be between .55 and .98 (Gur et al. 2010). Internal consistency was reported in a Dutch study to have a median Cronbach’s *α* of .86 across all Penn CNB subtests (Swagerman et al. 2016).

##### NIH Toolbox

Seven subtests from the NIH Toolbox for the Assessment of Neurological and Behavioral Function (http://www.nihtoolbox.org; Gershon et al. 2013; Heaton et al. 2014; Weintraub et al. 2013) were selected to assess intelligence. These were the Flanker Inhibitory Control and Attention Test (Flanker), the Dimensional Change Card Sort Test (CardSort), the List Sorting Working Memory Test (ListSort), the Picture Sequence Memory Test (PicSeq), the Oral Reading Recognition Test (ReadEng), the Picture Vocabulary Test (PicVocab), and the Pattern Comparison Processing Speed Test (ProcSpeed).

The Flanker is a test of executive function and measures ability to focus visual attention on a task-relevant stimulus while inhibiting attention towards task-irrelevant stimuli. During each trial, participants see an arrow in the center of a computer screen, which is flanked by similar stimuli on the left and right pointing in the same or the opposite direction as the target. The task is to indicate the direction of the central stimulus. There are 40 trials in total. The CardSort is a test of executive function as well but designed to measure cognitive flexibility. Participants are shown pictures that vary along two dimensions (e.g., shape and color). These must be assigned to one of two target pictures so that the pictures match either in shape or in color. The relevant criterion is displayed on a computer screen and varies without a predictable pattern among 40 trials. The ListSort measures working memory capacity. A series of stimuli is presented both visually, as objects on the computer screen, and orally, as spoken names. Participants are asked to repeat the stimuli in order of size. In the first condition, all stimuli come from the same category, but belong to one of two categories in the second condition. Participants first recite all stimuli from category one and then those from category two, always in order of size. The number of items increases with each trial and the task is discontinued as soon as a participant fails to complete two trials of the same length. The PicSeq is a test of episodic memory and measures acquisition, storage, and retrieval of new information. During each trial, pictures of objects and activities appear at the center of a computer screen. While the content of a single picture is described via an audio file, the picture moves across the screen so that there is a spatial arrangement of pictures at the end. Then all pictures are again randomly presented at the center of the screen and the participant must move them to match the spatial arrangement shown before. The ReadEng is a test of reading decoding skill. Participants are asked to pronounce letters and words shown on a computer screen as correctly as possible. Depending on performance, 30 to 40 items from a collection of 250 items are administered, while controlling frequency of word use, complexity of letter-sound relations, and orthographic typicality. The PicVocab measures general vocabulary knowledge. Participants hear a spoken word via an audio file and simultaneously see four different images (objects, actions, and/or depictions of concepts) on a computer screen. They choose the image that the spoken word labels. The ProcSpeed is a test of processing speed. Participants are asked to identify whether two images displayed side-by-side are identical or not. Within 90 seconds, as many pairs of images as possible should be evaluated (Gershon et al. 2013; Heaton et al. 2014; Weintraub et al. 2013).

The NIH Toolbox has been validated with several American samples (Heaton et al. 2014; Weintraub et al. 2013). For the subtests, Weintraub et al. (2013) reported test-retest reliabilities (intraclass correlation coefficients) between *r* = .78 and .99. Heaton et al. (2014) built and analyzed composite scores and found acceptable internal consistency (Cronbach’s α between .77 and .84) as well as excellent test-retest reliabilities between *r* = .86 and .92.

##### Acquisition of diffusion-weighted imaging data

All images were collected on a customized Siemens 3T Connectome Skyra scanner housed at Washington University in St. Louis, using a standard 32-channel Siemens head coil. Diffusion-weighted images were acquired using echo planar imaging (TR = 5520 ms, TE = 89.5 ms, flip angle = 78°, 111 slices, matrix size = 168 x 144, voxel size = 1.25 x 1.25 x 1.25 mm; Feinberg et al. 2010; Moeller et al. 2010; Setsompop et al. 2012; Xu et al. 2012). The complete diffusion-weighted imaging session was divided into six runs, each lasting approximately nine minutes and 50 seconds (total acquisition time of about one hour). The six runs represented three different gradient tables, once acquired in the right-to-left and in the left-to-right phase-encoding direction. Each gradient table comprised 90 diffusion weighting directions as well as six acquisitions with *b* = 0 s/mm^2^ interspersed throughout each run. Diffusion weighting was based on a multi-shell scheme consisting of equally distributed diffusion-weighted images for *b*-values of 1000, 2000, and 3000 s/mm^2^.

### Data set UMN

#### Participants

The UMN data set encompassed 335 participants (mean age: 26.3 years, SD = 5.0 years, 164 females) with sufficient data from intelligence testing to compute a general factor *g*. Since DWI data were not available for all participants, the final sample for the TBSS analysis was reduced to 251 participants (mean age: 26.2 years, SD = 4.9 years, 122 females). The age range was 20 to 40 years, therefore also representing young adulthood. Individuals who reported a history of neurologic or severe psychiatric disorders, current drug or alcohol problems, or current use of psychotropic medication (antipsychotics, anticonvulsants, and stimulants) were not admitted to the study. The study protocol was approved by the University of Minnesota Institutional Review Board and all participants gave written informed consent.

#### Measurement of intelligence

##### WAIS-IV

Intelligence was assessed using five subtests of the Wechsler Adult Intelligence Scale, fourth edition (WAIS-IV; Wechsler 2008): Block Design (WAIS_BD), Matrix Reasoning (WAIS_MR), Similarities (WAIS_SIM), Vocabulary (WAIS_VC), and Coding (WAIS_CD). The subtest WAIS_BD measures perceptual reasoning. Participants reproduce a two-dimensional pattern (shown on paper) with several red and white three-dimensional building blocks. Performance is quantified as the time needed to recreate the correct pattern. WAIS_MR is a subtest of perceptual reasoning as well. Participants are shown matrices composed of multiple elements, each arranged according to a specific rule. This rule must be identified to fill an empty spot within the matrix by choosing the correct piece among five options. The subtests WAIS_SIM and WAIS_VC assess verbal comprehension skills. During the WAIS_SIM, 18 word pairs are presented orally and participants describe qualitative similarity between the two words. The subtest WAIS_VC requires participants to define and/or describe up to 30 words or concepts. WAIS_CD is a test of processing speed. The numbers 1-9 are paired with different symbols. Participants are presented with a number sequence and add the corresponding symbols to as many numbers as possible within a given time limit.

The WAIS-IV subtests’ Cronbach’s *α*s have been reported to be between .84 and .94 and test-retest reliabilities to range between *r* = .69 and .91 (Wechsler 2008).

#### Acquisition of diffusion-weighted imaging data

All images were collected on a 3T Siemens Trio scanner at the Center for Magnetic Resonance Research (CMRR) at the University of Minnesota in Minneapolis, using a 12- channel head coil. Diffusion-weighted images were acquired using echo planar imaging (TR = 7900 ms, TE = 86 ms, flip angle = 90°, 64 slices, field of view = 2048 mm^2^, voxel size = 2 x 2 x 2 mm). Diffusion weighting was uniformly distributed along 71 directions. Nine measurements with a *b*-value of 1000 s/mm^2^ were conducted. The total acquisition time was 12 minutes, 34 seconds.

### Data set NKI

#### Participants

Data collection for the NKI sample is still ongoing. It is intended to investigate the neurologies of psychiatric disorders (Nooner et al. 2012). The “Enhanced Nathan Kline Institute - Rockland Sample” data set (Nooner et al. 2012) is part of the 1000 Functional Connectomes Project (http://fcon_1000.projects.nitrc.org) and we downloaded it from its official website (http://fcon_1000.projects.nitrc.org/indi/enhanced/). Since our study is focused on healthy participants, we included only individuals who did not report any history of psychiatric illness. Moreover, they also had to have complete intelligence test data. We used these to calculate the *g* factor (N = 417, mean age: 43.5 years, SD = 23.5 years, 273 females). For the final sample, usable for TBSS analysis, we had to exclude additional participants due to lack of DWI data (N = 396, mean age: 44.4 years, SD = 22.9 years, 259 females). Relative to the other data sets, which mainly contain data from young adults, the NKI sample had a much greater age range, 6-85 years. At 44.4 years, the mean age was also well above the means of the others (RUB: 27.3 years, HCP: 28.7 years, and UMN: 26.2 years). However, since exclusion of all participants outside the 20-40 range would have cost 306 participants, we included all participants with suitable data. The study protocol was approved by the Institutional Review Boards at the Nathan Kline Institute and Montclair State University. Written informed consent for the study was obtained from all participants or, for children, additionally from a legal guardian (Nooner et al., 2012).

#### Measurement of intelligence

The Wechsler Abbreviated Scale of Intelligence, second edition (WASI-II; Wechsler 2011), measured intelligence. The inventory has four subtests, Block Design (WASI_BD), Matrix Reasoning (WASI_MR), Similarities (WASI_SIM), and Vocabulary (WASI_VC), which are comparable to the subtests from the WAIS-IV (see Data set UMN). Despite being conceptually similar, the parallel subtests from the WAIS-IV and WASI-II include unique test items (McCrimmon and Smith 2012). WASI_BD has 13 items, WASI_MR 30 items, WASI_SIM 24 items, and WASI_VC 31 items. The WASI-II can be administered in about 30 minutes and is considered to be the measure of choice for brief intelligence assessments. Split-half reliabilities of the subtests varied between *r* = .87 and .91 in the child norming sample (6-16 years) and between *r* = .90 and .92 in the adult norming sample (17-90 years). Test-retest reliability was *r =* .79 in the child sample and .94 in the adult sample. The interrater reliabilities of the four subtests were between *r* = .94 and .99, considered exceptionally high (McCrimmon and Smith 2012).

#### Acquisition of diffusion-weighted imaging data

All images were collected on a Siemens Magnetom TrioTim syngo MR B17 scanner at the Nathan Kline Institute in Orangeburg, New York. Diffusion-weighted images were acquired using echo planar imaging (TR = 2400 ms, TE = 85 ms, flip angle = 90°, 64 slices, field of view = 212 mm, matrix size = 212 x 212, voxel size = 2 x 2 x 2 mm). Diffusion weighting was uniformly distributed along 128 directions using a *b*-value of 1500 s/mm^2^. In addition, nine volumes without diffusion weighting (*b* = 0 s/mm^2^) were obtained. The total acquisition time was five minutes, 58 seconds.

### Computation of the general factor of intelligence, *g*

Research on the psychometric structure of intelligence has modified and extended Spearman’s original ideas regarding the existence of *g*. In recent hierarchically organized models, *g* is placed at the apex of a hierarchy with broad cognitive domains at a lower level and narrow cognitive abilities at the basis (Flanagan and Dixon 2013; Schneider and McGrew 2012). There is considerable evidence for the existence of such structures, but their specifics depend on the tests and sample properties. Nonetheless, when ranges of tests included are broad, their *g* factors correlate for all practical purposes completely, e.g. Johnson et al. (2004); Johnson et al. (2008). Thus content of *g* is relatively unaffected by the tests from which it was generated, though the level of any one person’s factor score certainly could be. Measurement invariance does not hold across *g* ranges. For example, arithmetic tests tend to be processing speed tasks for people with high *g* levels but reasoning tasks for people with a low *g* levels. Furthermore, a person with average performance on various intelligence tests may have a standardized *g*-value that is below average in a highly intelligent sample and a *g*-value that is above average in a less intelligent sample. Since individuals’ *g* levels in different samples might differ from each other and because imaging data were obtained on different scanners (which also affects what is observed), it was not possible to combine the four data sets employed in our study.

We used the intelligence test scores of each data set to compute *g* factors for every participant. To do this, we regressed age, sex, age*sex, age^2^, and age^2^*sex from the test scores. We then developed a hierarchical factor model separately for each data set based on the standardized residuals, first using exploratory factor analysis, then confirmatory. We assessed model fit using Root Mean Square Error of Approximation (RMSEA), Standardized Root Mean Square Residual (SRMR), Comparative Fit Index (CFI), and Tucker-Lewis index (TLI). Values of RMSEA and SRMR less than .05 are considered good and values of CFI and TLI greater than .97 (Hu and Bentler 1999). We used these models to calculate regression-based *g-*factor scores for each participant, winsorizing outliers.

#### Confirmatory Factor Models

Figures 1 to 4 show the postulated factor models for the data sets, the z-standardized factor loadings, and the covariances between individual subtests. The confirmatory factor analyses of all data sets yielded quite good to excellent fit indices (see Table 1).

**Figure 1.**
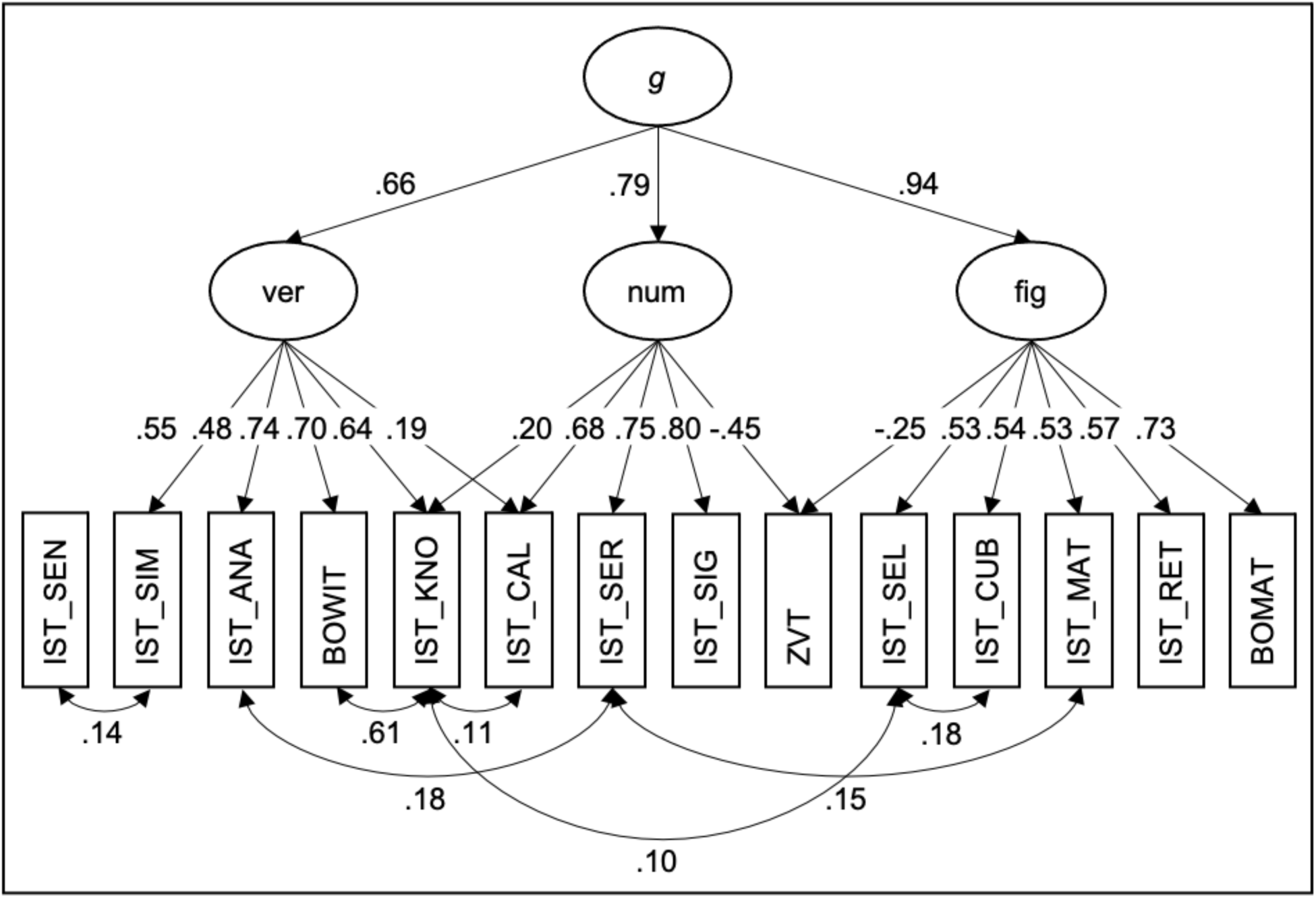
Factor structure of the RUB data set. *g* = general factor of intelligence, ver = verbal intelligence as broad cognitive domain, num = numerical intelligence as broad cognitive domain, fig = figural intelligence as broad cognitive domain, IST_SEN = subtest Sentence Completion of the I-S-T 2000 R, IST_SIM = subtest Similarities of the I-S-T 2000 R, IST_ANA = subtest Analogies of the I-S-T 2000 R, BOWIT = Bochumer Wissenstest, IST_KNO = parameter Knowledge of the I-S-T 2000 R, IST_CAL = subtest Calculations of the I-S-T 2000 R, IST_SER = subtest Number Series of the I-S-T 2000 R, IST_SIG = subtest Numerical Signs of the I-S-T 2000 R, ZVT = Zahlen-Verbindungs-Test, IST_SEL = subtest Figure Selection of the I-S-T 2000 R, IST_CUB = subtest Cubes of the I-S-T 2000 R, IST_MAT = subtest Matrices of the I-S-T 2000 R, IST_RET = parameter Retentiveness of the I-S-T 2000 R, BOMAT = Bochumer Matrizentest.

**Figure 2.**
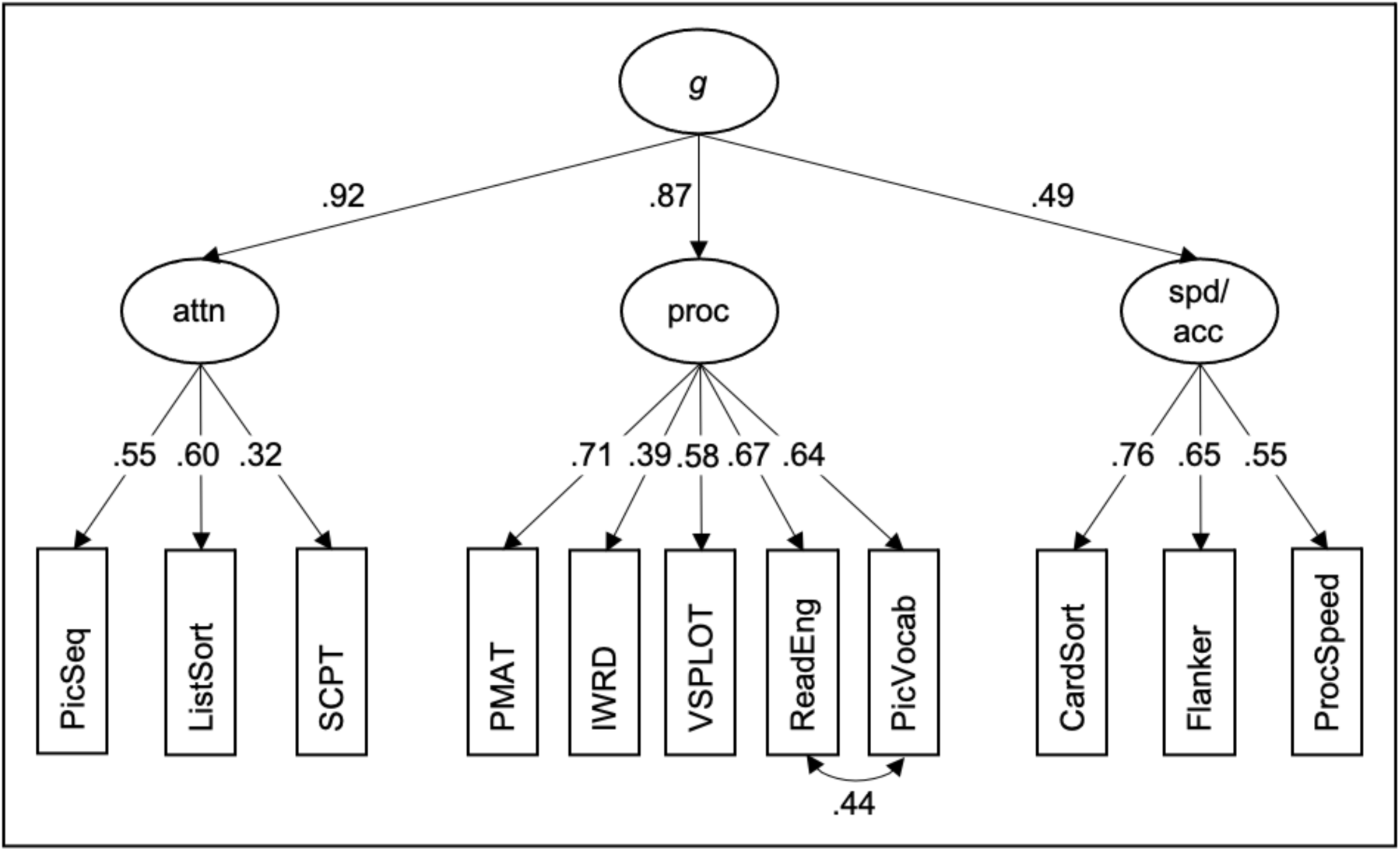
Factor structure of the HCP data set*. g* = general factor of intelligence, attn = attention as broad cognitive domain, proc = processing as broad cognitive domain, spd/acc = speed/accuracy as broad cognitive domain, PicSeq = subtest Picture Sequence Memory Test of the NIH Toolbox, ListSort = subtest List Sorting Working Memory Test of the NIH Toolbox, SCPT = subtest Short Penn Continuous Performance Test of the Penn CNB, PMAT = subtest Penn Matrix Reasoning Task of the Penn CNB, IWRD = subtest Penn Word Memory Test of the Penn CNB, VSPLOT = subtest Variable Short Penn Line Orientation Test of the Penn CNB, ReadEng = subtest Oral Reading Recognition Test of the NIH Toolbox, PicVocab = subtest Picture Vocabulary Test of the NIH Toolbox, CardSort = subtest Dimensional Change Card Sort Test of the NIH Toolbox, Flanker = subtest Flanker Inhibitory Control and Attention Test of the NIH Toolbox, ProcSpeed = subtest Pattern Comparison Processing Speed Test of the NIH Toolbox.

**Figure 3.**
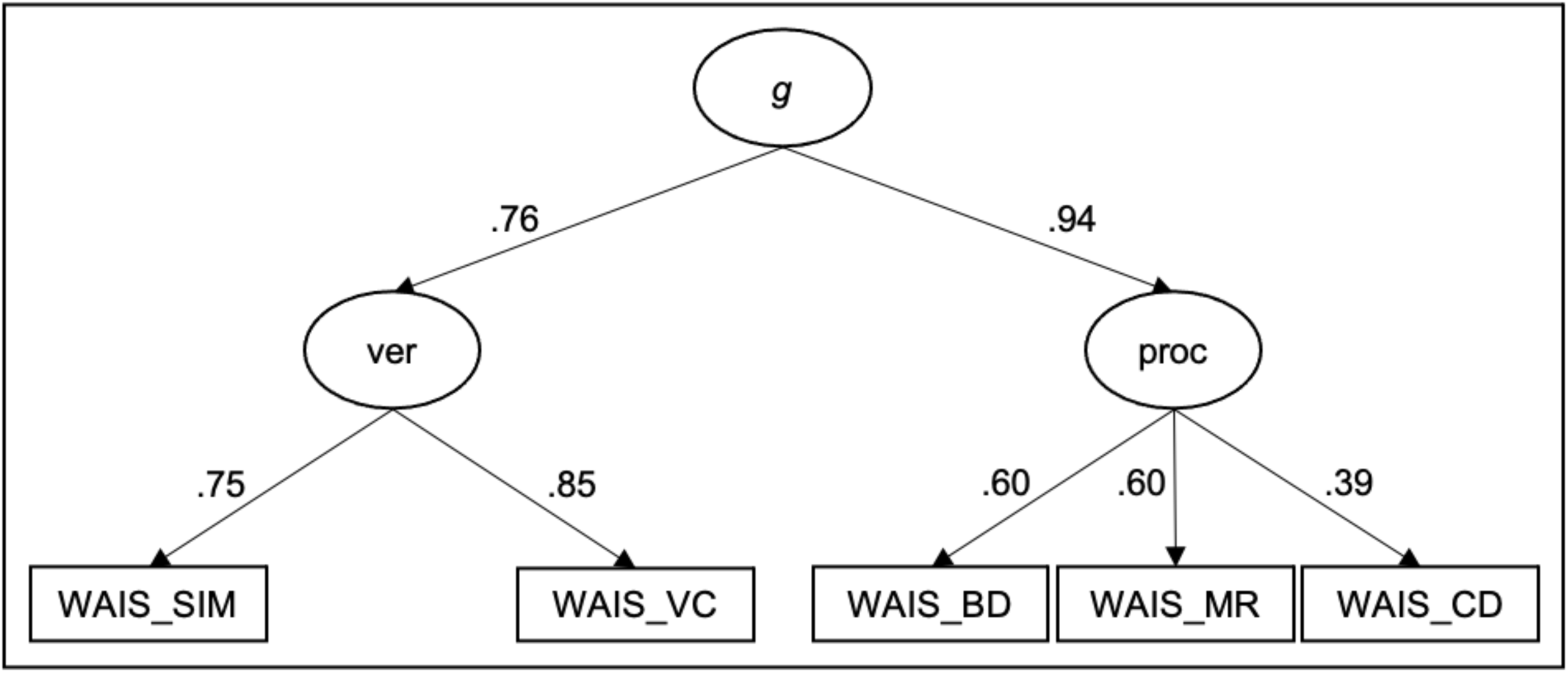
Factor structure of the UMN data set. *g* = general factor of intelligence. ver = verbal intelligence as broad cognitive domain, proc = processing as broad cognitive domain, WAIS_SIM = subtest Similarities of the WAIS-IV, WAIS_VC = subtest Vocabulary of the WAIS-IV, WAIS_BD = subtest Block Design of the WAIS-IV, WAIS_MR = subtest Matrix Reasoning of the WAIS-IV, WAIS_CD = subtest Coding of the WAIS-IV.

**Figure 4.**
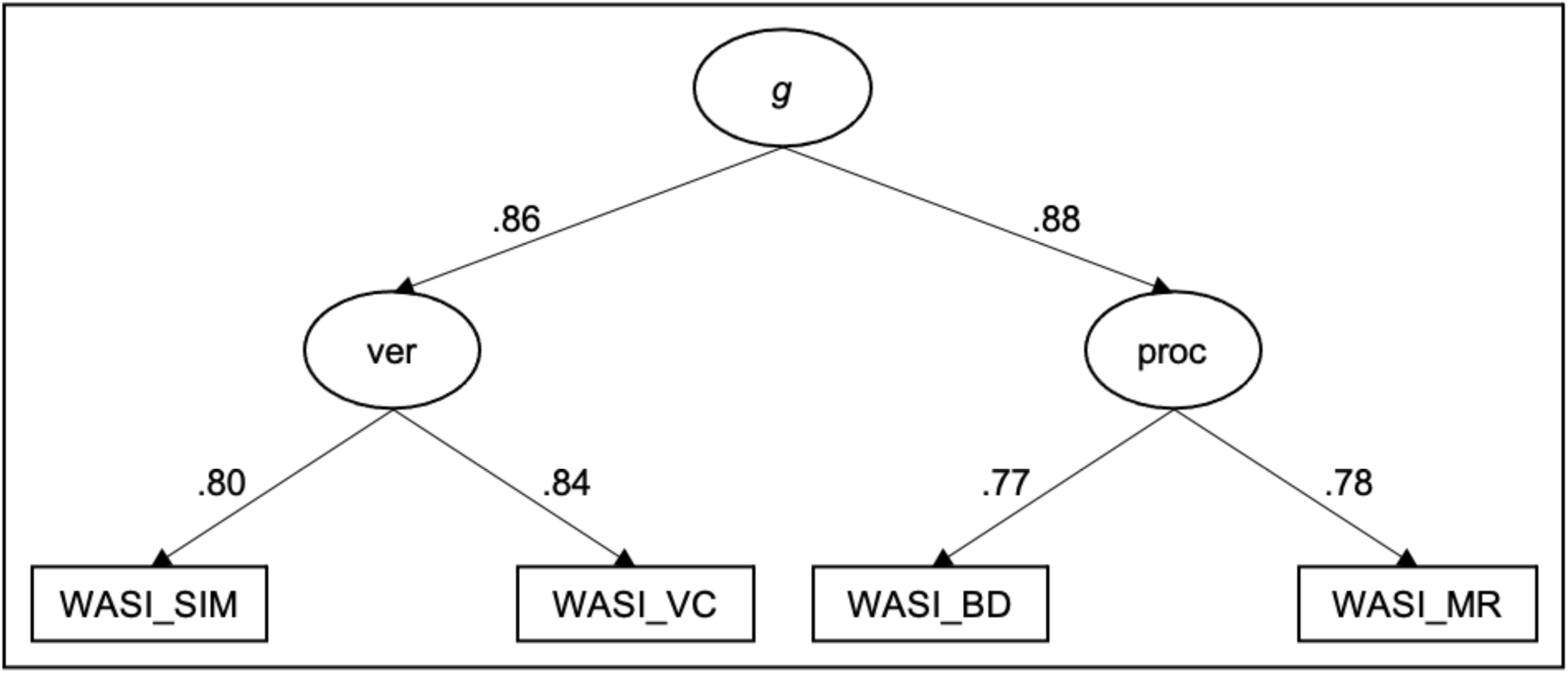
Factor structure of the NKI data set. *g* = general factor of intelligence, ver = verbal intelligence as broad cognitive domain, proc = processing as broad cognitive domain, WASI_SIM = subtest Similarities of the WASI-II, WASI_VC = subtest Vocabulary of the WASI-II, WASI_BD = subtest Block Design of the WASI-II, WASI_MR = subtest Matrix Reasoning of the WASI-II.

**Table 1.** Fit indices of the confirmatory factor analyses

| Model | $\chi^2$ | *df* | RMSEA | SRMR | CFI | TLI |
| --- | --- | --- | --- | --- | --- | --- |
| Data set RUB | 127.97*** | 64 | .042 | .033 | .979 | .969 |
| Data set HCP | 118.35*** | 40 | .041 | .028 | .973 | .963 |
| Data set UMN | 2.51 | 4 | .000 | .013 | 1.000 | 1.012 |
| Data set NKI | 0.13 | 1 | .000 | .002 | 1.000 | 1.009 |

RMSEA = Root Mean Square Error of Approximation. SRMR = Standardized Root Mean Square Residual. CFI = Comparative Fit Index. TLI = Tucker-Lewis-Index. \*\*\* $p < .001$ .

### Image processing and analysis

We processed and analyzed all data sets in the same manner. Since FA is one of the most commonly derived measures from diffusion data (Smith et al. 2006) and has been observed to be associated with intelligence in many studies (Genç and Fraenz 2021), we focused on FA. We used voxel-based statistical analysis of the FA data based on TBSS (Smith et al. 2006), which is part of Oxford Centre for Functional Magnetic Resonance Imaging of the Brain’s (FMRIB) Software Library (FSL), version 5.0.9 (Smith et al. 2004). First, DWI images were subjected to brain extraction using Brain Extraction Tool (BET; Smith 2002). Then, FA images were created by fitting tensor models to the raw diffusion data using FMRIB’s Diffusion Toolbox (FDT). We transformed the resulting FA images into a common space via FMRIB’s Nonlinear Image Registration Tool (FNIRT; Andersson et al. 2007a; 2007b), which uses b-spline representations of the registration warp fields (Rueckert et al. 1999). For this purpose, we chose the DTI template FSL_HCP1065_FA_1mm within FSL, which is based on 1065 participants from the Human Connectome Project and is available in Montreal Neurologic Institute (MNI) 152 standard space (1 x 1 x 1 mm). Next, we created and thinned mean FA images to generate mean FA skeletons representing the centers of all tracts common to the sample. We set the FA threshold at 0.20 to include only major white matter tracts and exclude peripheral tracts which are more vulnerable to intra- and inter-subject variability. Each participant’s aligned FA image was projected onto the skeleton by filling each skeleton voxel with the FA value of the nearest tract center. We used the resulting data to compute voxel-based cross-subject statistics.

### Statistical analysis

We used permutation-based inference (Nichols and Holmes 2002) to analyze voxel-based cross-subject image similarities. To this end, we executed the FSL tool “randomise” (Winkler et al. 2014) with 5,000 permutations for each analysis. Within the white matter skeleton of each data set, we used a general linear model (GLM) to identify positive and negative associations between *g* and FA while controlling age, sex, age*sex, age^2^, and age^2^*sex. We treated them as nuisance variables since they explain relatively little (∼10%) of the total variance in whole-brain average FA (Kochunov et al. 2015) and we were not interested in possible age and sex differences. We used threshold-free clustering enhancement (Smith and Nichols 2009) to avoid arbitrarily specifying a cluster-forming threshold a priori. We adjusted the resulting statistical parametric maps for multiple comparisons by the family-wise error rate thresholded at *p* < .05. We binarized them via the FSL tool “fslmaths”, so that voxels exhibiting a significant relation between *g* and FA were assigned 1 and all remaining voxels 0. We carried out each step separately in each data set.

As the focal final step, we compared our observations from the individual data sets to identify white matter areas exhibiting replicable structure-function associations. For this purpose, we used the FSL tool “fslmaths” to compute the sums of the four binarized maps depicting positive contrasts and the four binarized maps depicting negative contrasts (see Figure 5). This resulted in two statistical parametric maps with values between 0 (no positive/negative associations between *g* und FA in any data set) and 4 (positive/negative associations in all data sets). We thresholded these maps once again to generate conservative maps only showing those voxels that exhibited significant associations across all four data sets (100% consensus). We multiplied those conservative maps with thresholded (value 10) fiber tracts of the Johns Hopkins University White Matter Tractography Atlas, implemented in FSL, to determine the anatomical location of the voxels (Hua et al. 2008; Mori et al. 2005; Wakana et al. 2007). We averaged the FA values of all significant voxels within a voxel cluster for each participant. These mean FA values were related to *g* by calculating partial correlation with age, sex, age*sex, age^2^, and age^2^*sex as controls. We did this separately for each data set and results were visualized using scatter plots.

**Figure 5.**
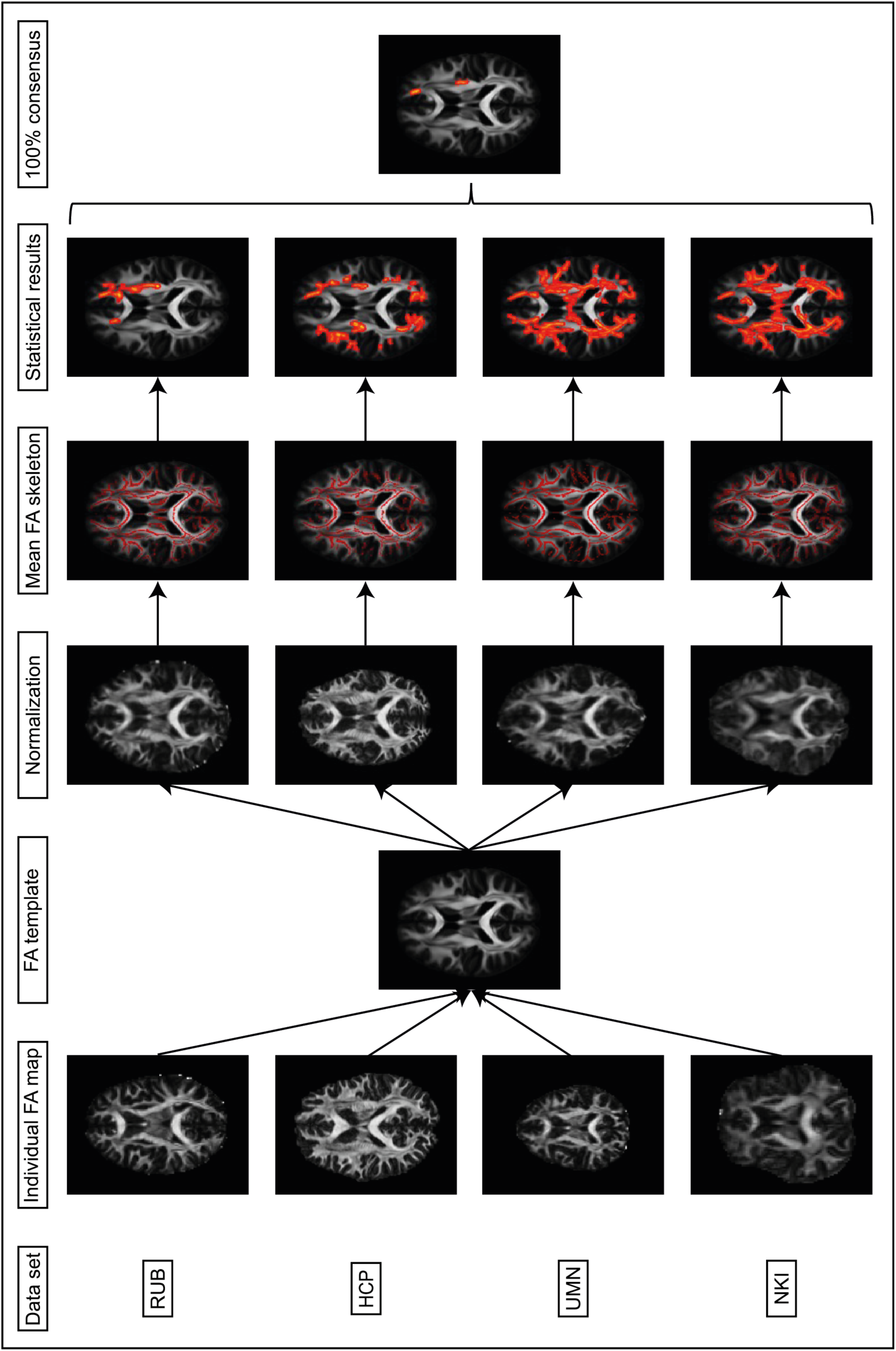
Methodological sequence depicting the different steps of the image analysis and statistical analysis. The TBSS approach was carried out for each data set separately. We used nonlinear registration to transform individual FA images to a common stereotactic space. By averaging all aligned images, we obtained mean FA maps (not shown). Next, we thinned these to generate white matter skeletons only including voxels at the center of fiber tracts common to all participants. We projected each participant’s aligned FA map onto a skeleton by filling the skeleton voxels with FA values from the nearest relevant tract center (not shown). We used the skeletonized FA maps to compute voxel-based cross-subject statistical comparisons. The second last column depicts statistical maps showing voxels that exhibited a significant positive relation between *g* and FA (controlled for age, sex, age*sex, age^2^, and age^2^*sex). The last image on the right shows voxels that matched across all four data sets.

### Additional exploratory analyses

We also took an exploratory and more liberal approach by creating brain maps including all voxels that exhibited significant associations in 3 out of 4 data sets (75% consensus).

Beyond that, we conducted further explorative analyses. These were based on previous studies’ reports that made different observations for broad, first-order intelligence factors such as verbal and performance abilities (Tamnes et al. 2010). First, we used each of the first-order intelligence factors from each data set (see Figures 1 to 4) as regressors on FA while adding age, sex, age*sex, age^2^, age^2^*sex, and the remaining first-order intelligence factors for each data set as nuisance factors. For example, the association between verbal intelligence and FA in the RUB data set was analyzed with age, sex, age*sex, age^2^, age^2^*sex, numerical intelligence, and figural intelligence serving as nuisance factors. Second, we removed the effects of *g* from all first-order intelligence factors and used these variables as regressors on FA, along with age, sex, age*sex, age^2^, and age^2^*sex as nuisance variables.

We also tried to compare the first-order intelligence factors by binarizing, adding, and thresholding their statistical parametric maps as described above for *g* to test whether there were robust observations among our four data sets below *g*. Since the factor models of our data sets had different first-order factors, it was not possible to compare them directly in all data sets. One example is the HCP data set which does not have a first-order intelligence factor related to only verbal abilities (see Figure 2). Nonetheless, we still tried to include this sample in our comparison of first-order intelligence factors. Hereby, we tested whether there was a robust relation between FA and verbal abilities by combining the results of the first-order intelligence factors ver (RUB, UMN, and NKI) and proc (HCP) (see Figures 1 to 4). For processing abilities, we combined the first-order intelligence factors fig (RUB) and proc (HCP, UMN, and NKI).

## Results

### Relations between *g* and FA

#### Main analysis with 100% consensus

In total 188 individual voxels, 0.12% of the white matter skeleton, exhibited significant positive associations between *g* and FA in all 4 data sets, controlling age, sex, age*sex, age^2^, and age^2^*sex. These voxels could be pooled into three contiguous clusters. Cluster “Forceps minor” was the largest and comprised 97 voxels. It overlapped completely with parts of the forceps minor as well as with crossing extensions of the anterior thalamic radiation, the cingulum-cingulate gyrus, and the inferior fronto-occipital fasciculus in the left hemisphere. Scatter plots illustrating the associations between this cluster’s mean FA and *g* are shown in Figure 6 (RUB: *r* = .15; HCP: *r* = .14; UMN: *r* = .13; NKI: *r* = .16). The second cluster “SLF” comprised 79 voxels and was located around the superior longitudinal fasciculus in the left hemisphere. Figure 7 shows the four scatter plots illustrating the associations between this cluster’s mean FA and *g* (RUB: *r* = .18; HCP: *r* = .14; UMN: *r* = .22; NKI: *r* = .12). The third cluster “Cingulum” was rather small and comprised 12 voxels. Since this cluster did not overlap with any of the thresholded fiber tracts, we used their unthresholded versions to assign the voxels to the fiber tracts. We observed matching voxels with fading extensions of the cingulum-cingulate gyrus, the inferior fronto-occipital fasciculus, and the anterior thalamic radiation in the left hemisphere. The four scatter plots illustrating the associations between this cluster’s mean FA and *g* are shown in Figure 8 (RUB: *r* = .14; HCP: *r* = .12; UMN: *r* = .13; NKI: *r* = .13). No voxels exhibited significant negative associations between *g* and FA in any of the 4 data sets.

**Figure 6.**
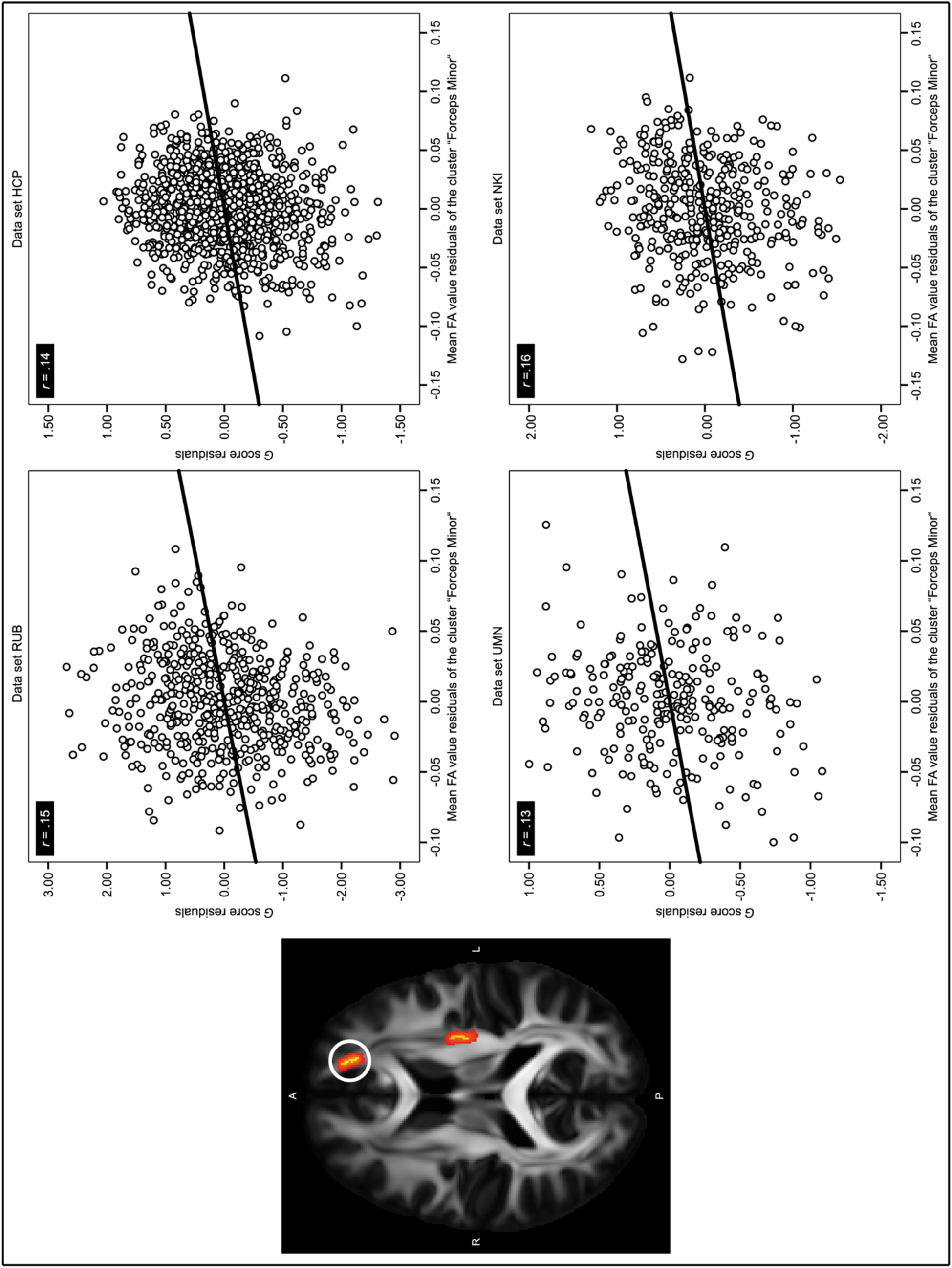
Associations between *g* and mean FA values from the cluster “Forceps minor”. The image on the left side shows the voxel cluster named “Forceps minor” (encircled). The FA values of these voxels were significantly positively associated with *g* in all four data sets (independent of effects of age, sex, age*sex, age^2^, and age^2^*sex). The voxels completely overlapped with parts of the forceps minor as well as with crossing extensions of the anterior thalamic radiation, the cingulum-cingulate gyrus, and the inferior fronto-occipital fasciculus in the left hemisphere. The right side of the figure shows four scatter plots, one for each data set. Here, mean FA values from cluster “Forceps minor” are plotted against standardized *g* values. Age, sex, age*sex, age^2^, and age^2^*sex were used as controlling variables.

**Figure 7.**
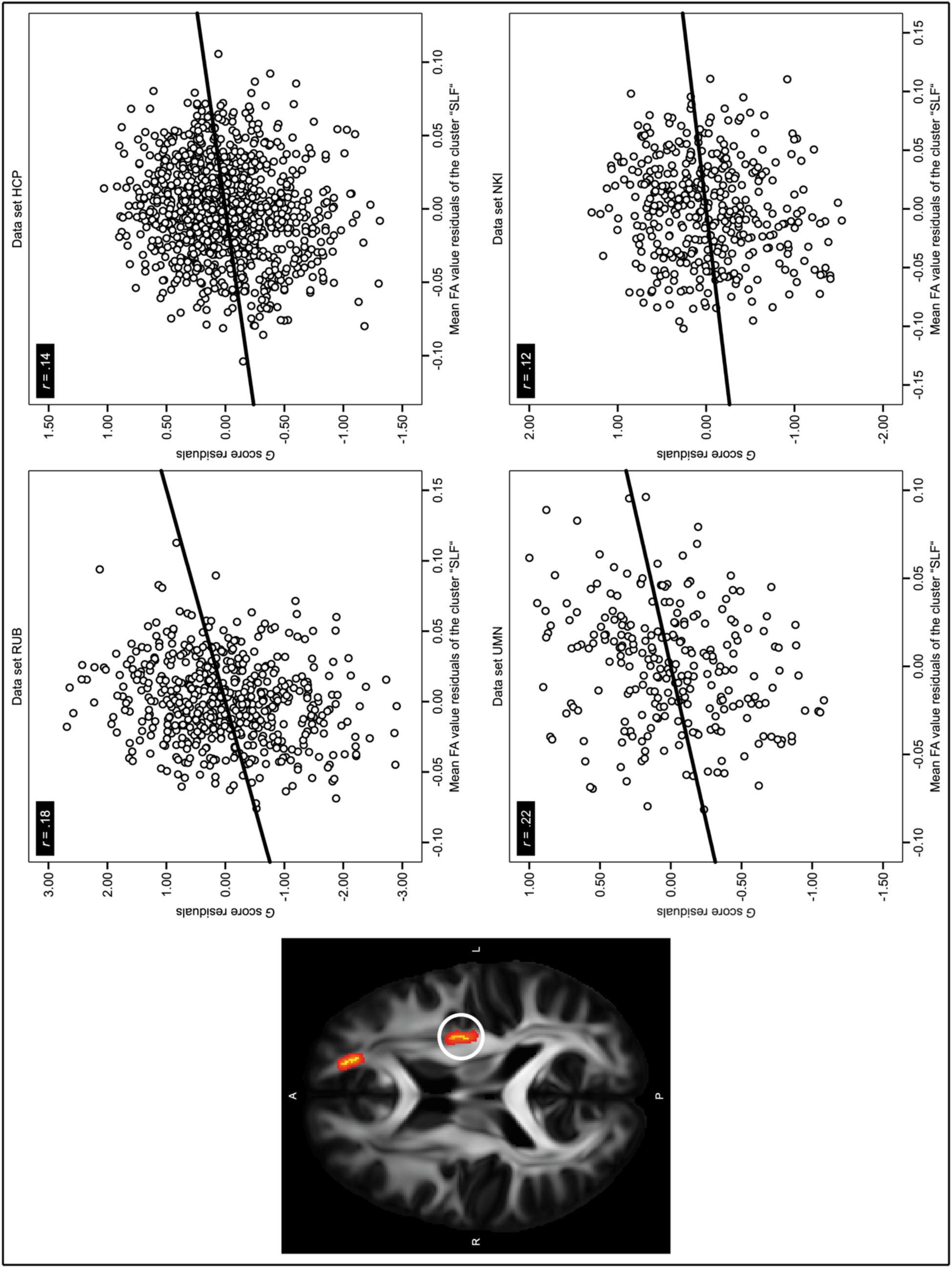
Associations between *g* and mean FA values from the cluster “SLF”. The image on the left side shows the voxel cluster named “SLF” (encircled). The FA values of these voxels were significantly positively associated with *g* in all four data sets (independent of the effects of age, sex, age*sex, age^2^, and age^2^*sex). The voxels were located around the superior longitudinal fasciculus in the left hemisphere. The right side of the figure shows four scatter plots, one for each data set. Here, mean FA values from cluster “SLF” are plotted against standardized *g* values. Age, sex, age*sex, age^2^, and age^2^*sex were used as controlling variables.

**Figure 8.**
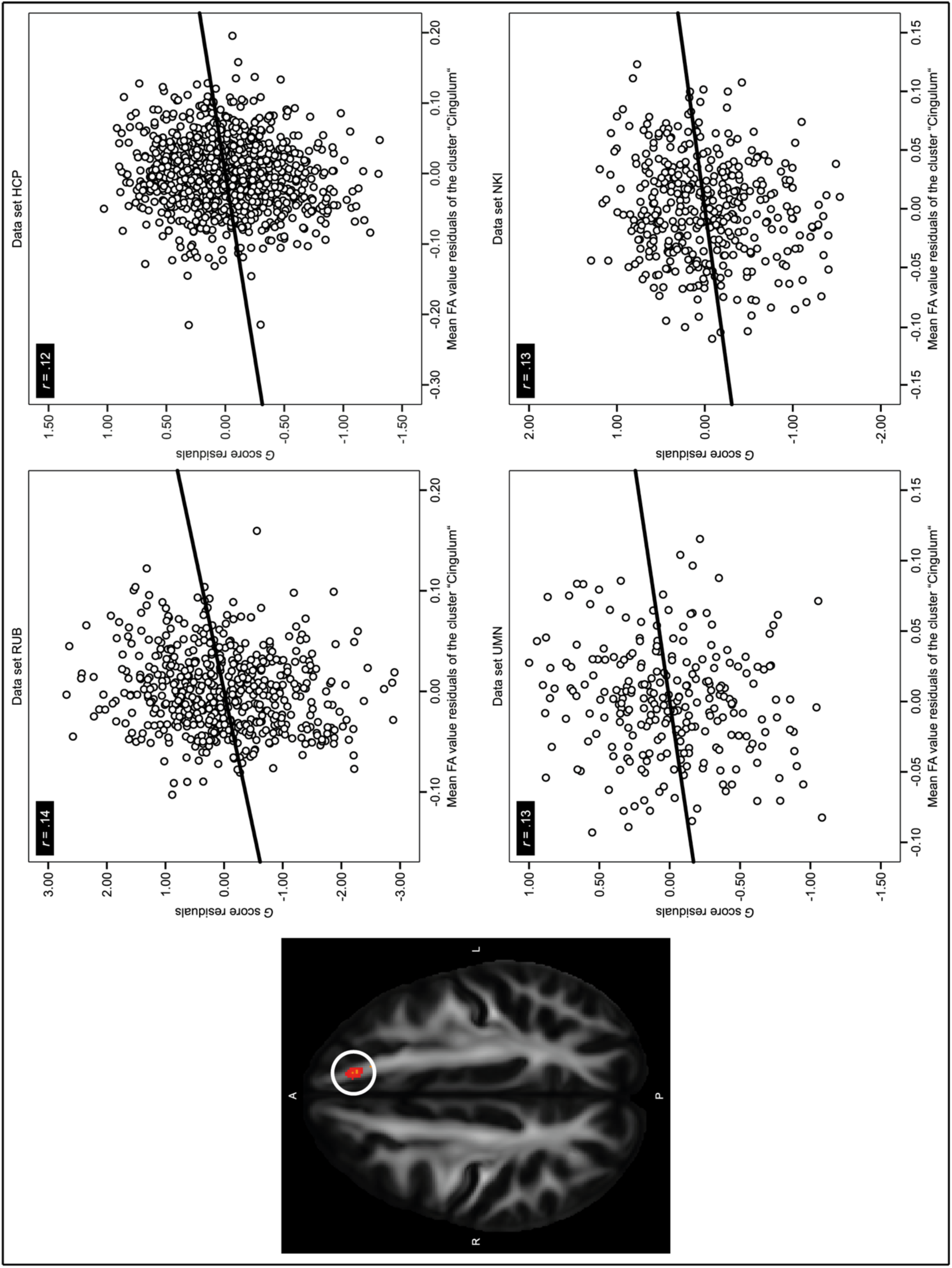
Associations between *g* and mean FA values from the cluster “Cingulum”. The image on the left side shows the voxel cluster named “Cingulum” (encircled). The FA values of these voxels were significantly positively associated with *g* in all four data sets (independent of the effects of age, sex, age*sex, age^2^, and age^2^*sex). The voxels overlapped with fading extensions of the unthresholded fiber tracts cingulum-cingulate gyrus, inferior fronto-occipital fasciculus, and anterior thalamic radiation in the left hemisphere. The right side of the figure shows four scatter plots, one for each data set. Here, mean FA values from cluster “Cingulum” are plotted against standardized *g* values. Age, sex, age*sex, age^2^, and age^2^*sex were used as controlling variables.

#### Exploratory approach with 75% consensus

The more liberal approach, requiring results to replicate in 3 of the 4 data sets, yielded 8364 voxels, 5.5% of the white matter skeleton, with significant positive associations between *g* and FA, controlling age, sex, age*sex, age^2^, and age^2^*sex. As depicted in Supplemental Figure 1, these voxels were widely scattered across the skeleton. Table S1 shows the distribution of significant voxels in relation to various major white matter fiber tracts.

### Exploratory approach for first-order intelligent factors below *g*

As mentioned above, we also tested whether there were robust associations below the level of *g*. The different analyses focused on first-order intelligence factors did not yield consistent results for 100% consensus, 75% consensus, or 50% consensus. Hence, we do not present our observations of single data sets.

## Discussion

Previous research focused on the relations between general intelligence and white matter microstructure in healthy participants has yielded mixed results. Hence, the primary goal of this study was to find replicable structure-function associations between general intelligence and white matter FA. Indeed, our analyses, involving a TBSS approach across four large, independent, cross-sectional samples, led to the conclusion that such replicable associations exist. We were able to identify a total of 188 voxels, 0.12% of the white matter skeleton, that exhibited significant positive relations between *g* and FA across all four data sets, controlling age, sex, age*sex, age^2^, and age^2^*sex. These voxels formed three contiguous clusters. The first was located around the forceps minor, crossing with extensions of the anterior thalamic radiation, the cingulum-cingulate gyrus, and the inferior fronto-occipital fasciculus in the left hemisphere. The second was located around the left-hemispheric superior longitudinal fasciculus. The third was located around the left-hemispheric cingulum-cingulate gyrus, crossing with extensions of the anterior thalamic radiation and the inferior fronto-occipital fasciculus.

There were no voxels exhibiting significant negative associations between *g* and FA in any of the four data sets. In this regard, our observations were consistent with previous research. Multiple studies have examined the associations between various measures of intelligence and FA by means of different approaches such as ROI-based, tract-based, whole-brain-based, or TBSS-based analyses. Despite these differences in design, these studies almost exclusively reported positive associations (Genç and Fraenz 2021). This suggests that individuals with higher intelligence scores tend to have greater degrees of linear white matter organization. However, the exact neurobiological underpinnings driving FA signal differences remain unclear. Hence, future studies are needed to examine the specific ways in which FA is affected by various factors like axon diameter, fiber density, myelin concentration, or the distribution of fiber orientation (Beaulieu 2002; Friedrich et al. 2020; Jones et al. 2013; Le Bihan 2003). It remains to be seen whether it will be possible to find universal neurobiological underpinnings. Since the way the brain gets wired in infancy and throughout further life varies among individuals, due to intertwined genetic and experiential differences, there is a possibility that there are basically no universal factors (Hasson et al. 2020).

Not only do our observations match previous research in direction of correlations, but the loci of voxels we were able to identify are consistent with those reported by other studies. Voxels were situated in regions of the forceps minor, the anterior thalamic radiation, the cingulum-cingulate gyrus, the inferior fronto-occipital fasciculus, and the superior longitudinal fasciculus in the left hemisphere. All these fiber tracts have been reported in previous TBSS-studies, albeit not consistently (Dunst et al. 2014; Malpas et al. 2016; Tamnes et al. 2010).

Fibers running through the genu, i.e. the anterior part of the corpus callosum, form the forceps minor (Catani and Thiebaut de Schotten 2008). As summarized by Genç and Fraenz (2021), the genu of the corpus callosum is the brain region in which FA is most often associated with interindividual differences in intelligence. In addition to FA, other callosal characteristics like cross-sectional area, density, or thickness have been related to intelligence (Atkinson et al. 1996; Hulshoff Pol et al. 2006; Luders et al. 2007), which underlines the relevance of this fiber tract for intellectual performance. The corpus callosum is the largest commissural fiber bundle in the brain and consists of approximately 200 million axons (Aboitiz et al. 1992). It connects the left and the right hemispheres and is thus crucial for interhemispheric transfer (van der Knaap and van der Ham 2011). While the left hemisphere (in most people) predominantly mediates processing of letters, words, language-related sounds, complex voluntary movements, verbal memory, speech, reading, writing, and arithmetic, the right hemisphere is predominantly crucial for processing of complex geometric patterns, faces, nonlanguage environmental sounds, music, tactile recognition of complex system patterns, movements in spatial patterns, nonverbal memory, prosody, geometry, sense of direction, and mental rotation of shapes (Kolb and Whishaw 2015). Besides lateralized functions, the two hemispheres have different roles in inferential reasoning (Marinsek et al. 2014). The left hemisphere seems to strive to reduce uncertainty by creating explanations, making inferences and bridging gaps in information, whereas the right hemisphere strives to resolve inconsistency by detecting conflicts, updating beliefs, supporting mental set-shifts, and monitoring and inhibiting behavior (Marinsek et al. 2014), but it is also the primary location of abstract reasoning processes such as those involved in analogies and metaphors (Diaz and Eppes 2018). For intelligent behavior in real life just as for good performance in artificial tasks used in intelligence tests, it seems essential that both hemispheres’ specialized functions and inferential reasoning strategies are important.

Given this, it is not surprising that agenesis of the corpus callosum, a congenital condition describing the absence of all or part of the corpus callosum, is associated with reduced interhemispheric transfer of sensory-motor information, reduced speed of cognitive processing, as well as deficits in complex reasoning and novel problem-solving (Brown and Paul 2019). This, in turn, can impact a wide range of cognitive and psychosocial functions (Brown and Paul 2019; Paul et al. 2016; Siffredi et al. 2013; Siffredi et al. 2018). Fibers of the genu link the prefrontal cortices across the hemispheres (Catani and Thiebaut de Schotten 2008). Macrostructural and functional properties of the prefrontal cortex have been repeatedly associated with intelligence (Basten et al. 2015; Deary et al. 2010a; Jung and Haier 2007). In general, the prefrontal cortex is highly relevant for higher cognitive skills such as abstract reasoning, problem solving, memory retrieval, attention, working memory, social interactions, language, and planning (Cabeza and Nyberg 2000; Wood and Grafman 2003). Furthermore, our observation that FA within the genu was associated with general intelligence is also supported by a recent functional connectivity study by Fraenz et al. (2021). This study examined the association between fluid intelligence and functional resting-state connectivity. They identified five connections exhibiting robust and replicable relations, including two interhemispheric connections in the frontal cortex (Fraenz et al. 2021).

The anterior thalamic radiation is a projection tract that connects the thalamus to the frontal lobe (Mori et al. 2002; Mori et al. 2005). Of all subcortical structures, thalamus volume seems to be most strongly associated with interindividual differences in intelligence (Bohlken et al. 2014; Cox et al. 2019). In addition, the thalamus has a complex connectivity profile, and its various nuclei establish connections to many areas of the brain (Aggleton et al. 2010; Behrens et al. 2003). Although the thalamus has traditionally been considered to serve merely as a relay station for cortical inputs, more recent observations suggest that its role in cognition could be much broader. It is conceivable that the thalamus also performs dynamic computations that take contextual information into account and reconfigure cortical representations (Dehghani and Wimmer 2019; Rikhye et al. 2018).

The cingulum is a medial associative fiber bundle that runs within the cingulated gyrus from the orbital frontal regions along the dorsal surface of the corpus callosum down towards the temporal lobe (Bubb et al. 2018; Catani and Thiebaut de Schotten 2008). Its fibers form intracortical connections between the medial frontal, parietal, occipital, and temporal lobes as well as different portions of the cingulated cortex. The fiber bundle is also part of the limbic system and one component of the Papez circuit (Papez 1937) constituting connections among the anterior thalamic nuclei, the parahippocampal region, and the cingulate cortex (Buyanova and Arsalidou 2021; Catani and Thiebaut de Schotten 2008). The cingulum appears to be involved in various cognitive domains such as cognitive control, attention, executive functions, memory, language, and visual-spatial functions (Bettcher et al. 2016; Bubb et al. 2018; Buyanova and Arsalidou 2021; Kantarci et al. 2011; Takahashi et al. 2010).

The inferior fronto-occipital fasciculus forms a major association fiber bundle linking the orbitofrontal cortex with the ventral occipital lobe (Catani and Thiebaut de Schotten 2008). Studies suggest that the inferior fronto-occipital fasciculus participates in semantic and visual processing as well as attention (Buyanova and Arsalidou 2021; Catani and Thiebaut de Schotten 2008; Leng et al. 2016).

The superior longitudinal fasciculus is a major white matter tract that connects frontal and opercular areas with the temporoparietal junction and parietal regions (Buyanova and Arsalidou 2021), allowing widespread intracortical information exchange. It is a matter of debate whether the arcuate fasciculus, which connects brain areas relevant for language processing (Broca’s and Wernicke’s area), can be considered part of the superior longitudinal fasciculus or is merely adjacent to it (Cox et al. 2019; Dick and Tremblay 2012; Kamali et al. 2014). Buyanova and Arsalidou (2021) noted that the right superior longitudinal fasciculus has been associated with cognitive functions such as attention (Frye et al. 2010) and visuospatial abilities (Hoeft et al. 2007), whereas the left superior longitudinal fasciculus has been observed to be crucial for language (Dick and Tremblay 2012) and reading skills (Frye et al. 2010). Buyanova and Arsalidou (2021) further stated that the arcuate fasciculus has been related to reasoning abilities and language processing (Lebel and Beaulieu 2009; Zemmoura et al. 2015). Therefore, both fiber tracts seem to be crucial for higher-order language functions (Friederici 2009). Language, in turn, is viewed as an important cognitive tool for problem solving since the lexicon symbols encapsulate abstract notions, making them more readily manipulable (Varley 2007). Grammatical mechanisms have similar roles in articulating relations among entities. Hence, language in the form of inner speech may allow tasks to be broken into finite series of sub-steps that guide reasoning processes (Varley 2007). Based on this inference, it is not surprising that the superior longitudinal fasciculus is one of the four fiber tracts being most often associated in the kinds of tasks used in intelligence tests (Genç and Fraenz 2021), especially given the constraints (e.g. many, extremely finite, rigidly structured items, administration under tight time and space conditions) involved in attempting to measure intelligence.

Our observations suggest that these brain regions play a vital roles in intelligence test performance via white matter tract integrity and organization, which is supported by previous research. Jung and Haier (2007) also posited the relevance of these fiber tracts in their P-FIT model. They proposed that working on intelligence test reasoning tasks involves multiple processing stages and harmonic interplay of the brain regions constituting their ‘P-FIT’ network. More precisely, they suggested that brain regions in the temporal and occipital lobes are crucial in successfully recognizing and initially processing sensory information. Subsequently, they presumed that the parietal cortex is essential for the interpretation, abstraction, and elaboration of the information’s symbolic content. The parietal cortex is believed to interact with frontal regions, which are thought to orchestrate generation and testing of potential solutions to given problems. Once a solution has been selected, it is thought that the anterior cingulate cortex chooses an appropriate reaction and inhibits alternative responses. Based on this, Jung & Haier (2007) proposed that the rapid and error-free transfer of information from posterior to frontal brain areas depends on underlying white matter integrity. They also emphasized the importance of information exchange between parietal and frontal association areas, which would highlight a role for the superior longitudinal fasciculus (Jung and Haier 2007). Therefore, our observations relating the superior longitudinal fasciculus to general intelligence supported the P-FIT model. Our cingulum observations fit within the P-FIT network. As noted by Fraenz et al. (2021), the P-FIT network is not organized exclusively intra-hemispherically. Hence, interhemispheric information transfer between prefrontal areas, e.g. via the forceps minor, seems to be consistent as well.

The P-FIT model does not propose direct connections between occipital and (orbito-)frontal areas. However, our observations, highlighting the importance of the inferior fronto-occipital fasciculus, did not necessarily contradict the model, given that this fiber tract also connects distal cortical regions of the P-FIT network. Instead, additional connections offer the possibility of more parallel flows of information. Since individuals who score identically in an intelligent test may use different cognitive strategies as well as different brain structures to reach their performance level (Deary et al. 2010a), there may be more than one adequate solution path and overall good brain function may be more important for general intelligence than using any specific parts well. Furthermore, we only observed significant voxel associations between FA and general intelligence in the left hemisphere, which is more relevant for language-related processing (Kolb and Whishaw 2015) as well as reasoning that strives to reduce uncertainty (Marinsek et al. 2014). Again, this is consistent with the P-FIT model given that half the Brodmann areas included in the P-FIT model predominantly exhibited left-hemispheric associations with intelligence, but as mentioned above this might be related to the constraints involved in attempting to measure intelligence.

Jung and Haier (2007) assumed that brain regions beyond the cerebral cortex, such as thalamus, hippocampus, and cerebellum, are involved only in rather basic functions. Hence, they believed that they would not contribute to interindividual intelligence differences significantly. However, more recent studies indicate that the thalamus and the hippocampus as well as their connections could play more important roles in reasoning than originally thought (Bohlken et al. 2014; Cox et al. 2019; Deary et al. 2022; Dehghani and Wimmer 2019; Rikhye et al. 2018). Our observations, involving the anterior thalamic radiation, supported these studies in suggesting that the P-FIT model (Jung and Haier 2007) needs some updating, which is only to be expected after 15 years more research.

We initially took a rather conservative analytical approach. To be considered for discussion, voxels had to exhibit significant associations between *g* and FA across all four data sets (100% consensus). A more liberal threshold (75% consensus) yielded about 44 times more voxels. Moreover, as illustrated in Supplemental Figure 1, significant voxel clusters were no longer exclusively located in the left hemisphere. However, Table S1 indicates that most significant voxels could be assigned to fiber tracts in the left hemisphere. As outlined above, the left and the right hemisphere differ in their specialized functions (Kolb and Whishaw 2015; Marinsek et al. 2014), which emphasizes the relevance of both hemispheres and their functional interaction for intelligent performances. One possible factor of influence regarding the found dominance of the left hemisphere could be language lateralization since language in the form of inner speech is considered as an important cognitive tool for problem solving (Varley 2007), especially given the constraints involved in intelligence tests, all intelligence test tasks rely on language to be executed (if only to comprehend the instructions for completing the tasks) and most people show a left hemisphere dominance of language lateralization accompanied by structural hemispheric asymmetries (Mazoyer et al. 2014; Ocklenburg and Güntürkün 2018a; 2018b).

## Limitations

Making use of multiple samples, as we did is more likely to yield replicable observations. However, the question arises why particular observations in one sample failed to replicate in other data sets. This might be because there is no robust association between *g* and FA, but it might also be that there are real effects that were simply too small to be detected as significant in the smaller samples, though they could be detected in the bigger ones. Another possible explanation might stem from differences among data sets. The four data sets included in our study used different intelligence tests, had different sample sizes, sex ratios, age distributions, and image acquisition protocols and came from different populations that they represented to different degrees. For example, the sample RUB mainly consisted of German university students who are not representative of the European population in age, educational background, or ethnic composition. Therefore, one should not draw conclusions about humans in general population based on our results. We also had no direct way to assess to what degrees our samples had similar mean levels of intelligence and *g* scores. We attempted to minimize the effects of these differences by calculating *g* factor scores, standardizing data processing for all data sets, and statistically controlling age, sex, age*sex, age^2^, and age^2^*sex.

Nevertheless, these differences might have hindered detection of potential associations and/or distorted those we did observe. In particular, our exploratory analyses of different first-order intelligence factors did not lead to robust results which could be replicated in all four data sets. Our observations were not consistent with Tamnes et al. (2010), who reported significant positive associations between FA and verbal/performance abilities. This could be because our analyses of these narrower intelligence factors controlled *g* itself, which theirs did not, thus examining only factor specific-variance. *g* typically explains about 40% of total variance in typical test batteries (Deary et al. 2010a). To resolve such inconsistencies, future studies should focus on specific intelligence factors, though keeping in mind that no factor identified in the manners used actually ‘carves nature at its joints’. They all vary considerably depending specific test battery content and sampling. In general, use of complementary methods, including fine-grained cortical parcellation schemes in combination with diffusion-weighted imaging and graph theory, may lead to new insights and are highly encouraged.

## Conclusion

In conclusion, we reported replicable associations between general intelligence and FA among four different cross-sectional data sets. By analyzing data from more than 2000 healthy participants, we were able to observe a total of 188 voxels with significant positive associations between *g* and FA in all four data sets, controlling age, sex, age*sex, age^2^, and age^2^*sex. These voxels were located around the forceps minor, crossing with extensions of the anterior thalamic radiation, the cingulum-cingulate gyrus, and the inferior fronto-occipital fasciculus in the left hemisphere, around the left-hemispheric superior longitudinal fasciculus, and around the left-hemispheric cingulum-cingulate gyrus, crossing with extensions of the anterior thalamic radiation and the inferior fronto-occipital fasciculus. Our observations do not imply that other brain’s white matter areas are irrelevant for intellectual performance, only that the mentioned fiber tracts show relatively stronger associations with intelligence than others. For the most part, our observations were consistent with previous research on the associations between white matter correlates and intelligence differences. We hope that future studies will make use of multiple samples because it is more likely to avoid false positive observations and could ultimately yield truly robust findings.

## Funding

This work was supported by the Deutsche Forschungsgemeinschaft (GU 227/16-1).

## Acknowledgments

The authors thank all research assistants for their support during the behavioral measurements. Further, the authors thank PHILIPS Germany (Burkhard Mädler) for the scientific support with the MRI measurements as well as Tobias Otto for technical assistance.

Address correspondence to Erhan Genç, Department of Psychology and Neuroscience, Leibniz Research Centre for Working Environment and Human Factors (IfADo), Ardeystraße 67, 44139 Dortmund, Germany.

## Notes

### Competing Interest Statement

The authors have declared no competing interest.

