## Supplementary Material for "Robust associations between white matter microstructure and general intelligence"

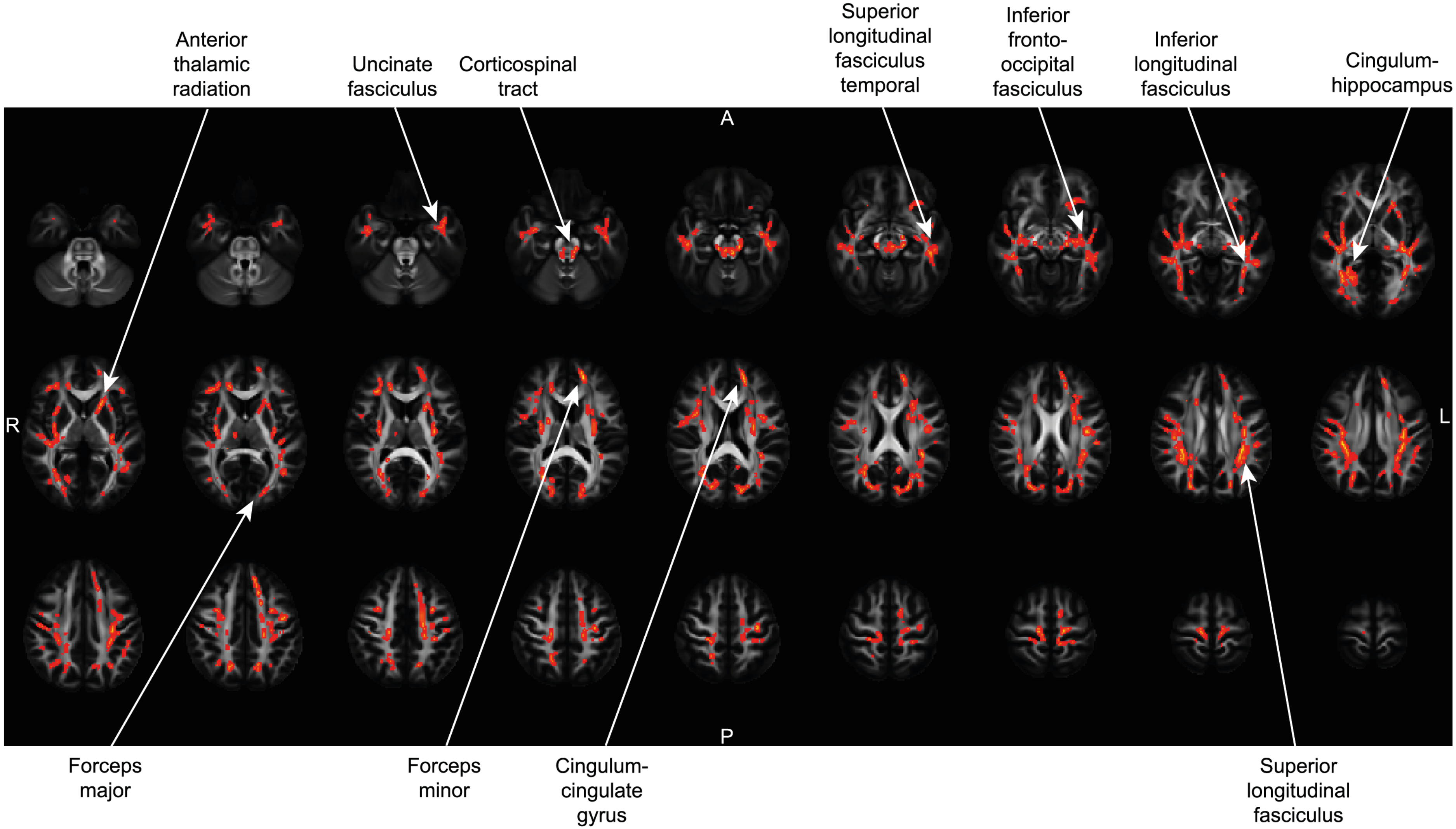
Anterior thalamic radiation
Uncinate fasciculus
Corticospinal tract
Superior longitudinal fasciculus temporal
Inferior fronto-occipital fasciculus
Inferior longitudinal fasciculus
Cingulum-hippocampus
A
R
L
P
Forceps major
Forceps minor
Cingulum-cingulate gyrus
Superior longitudinal fasciculus

**Supplemental Figure 1**. Voxels exhibiting significant positive associations between *g* and FA in 75% of data sets. Brain slices depict an exemplary selection of voxels showing significant positive relations between *g* and FA (controlled for the effects of age, sex, age*sex, $age^2$, and $age^2$*sex) in three out of the four data sets (75% consensus). Each arrow points at a voxel cluster that overlaps with a major white matter fiber tract. Apart from the Cingulum-hippocampus fiber tract, all fiber tracts are shown in the left hemisphere.

**Table S1** Distribution of significant voxels across white matter fiber tracts

| JHU White Matter Tract | Number of Voxels |
|---|---|
| Anterior thalamic radiation Left | 266 |
| Anterior thalamic radiation Right | 70 |
| Cingulum (Cingulate gyrus) Left | 27 |
| Cingulum (Cingulate gyrus) Right | 3 |
| Cingulum (Hippocampus) Left | 0 |
| Cingulum (Hippocampus) Right | 7 |
| Corticospinal tract Left | 210 |
| Corticospinal tract Right | 128 |
| Forceps major | 311 |
| Forceps minor | 420 |
| Inferior fronto-occipital fasciculus Left | 474 |
| Inferior fronto-occipital fasciculus Right | 583 |
| Inferior longitudinal fasciculus Left | 500 |
| Inferior longitudinal fasciculus Right | 404 |
| Superior longitudinal fasciculus Left | 974 |
| Superior longitudinal fasciculus Right | 651 |
| Superior longitudinal fasciculus (temporal) Left | 593 |
| Superior longitudinal fasciculus (temporal) Right | 313 |
| Uncinate fasciculus Left | 170 |
| Uncinate fasciculus Right | 49 |

The table summarizes the distribution of voxels, exhibiting significant positive associations between *g* and FA in three out of the four data sets, across various major white matter tracts (thresholded with the value 10) from the Johns Hopkins University White Matter Tractography Atlas (Hua et al., 2008; Mori et al., 2005; Wakana et al., 2007). Please note that only 6153 of the 8364 voxels (73.57%) could be assigned to any of the white matter tracts. Moreover, a single voxel might also overlap with more than one fiber tract. Hence, this table is only intended to outline a rough distribution of the significant voxels.
